# Antigen spacing on protein nanoparticles influences antibody responses to vaccination

**DOI:** 10.1101/2023.05.23.541980

**Authors:** Daniel Ellis, Annie Dosey, Seyhan Boyoglu-Barnum, Young-Jun Park, Rebecca Gillespie, Hubza Syeda, Yaroslav Tsybovsky, Michael Murphy, Deleah Pettie, Nick Matheson, Sidney Chan, George Ueda, Jorge A. Fallas, Lauren Carter, Barney S. Graham, David Veesler, Masaru Kanekiyo, Neil P. King

## Abstract

Immunogen design approaches aim to control the specificity and quality of antibody responses to enable the creation of next-generation vaccines with improved potency and breadth. However, our understanding of the relationship between immunogen structure and immunogenicity is limited. Here we use computational protein design to generate a self-assembling nanoparticle vaccine platform based on the head domain of influenza hemagglutinin (HA) that enables precise control of antigen conformation, flexibility, and spacing on the nanoparticle exterior. Domain-based HA head antigens were presented either as monomers or in a native-like closed trimeric conformation that prevents exposure of trimer interface epitopes. These antigens were connected to the underlying nanoparticle by a rigid linker that was modularly extended to precisely control antigen spacing. We found that nanoparticle immunogens with decreased spacing between closed trimeric head antigens elicited antibodies with improved hemagglutination inhibition (HAI) and neutralization potency as well as binding breadth across diverse HAs within a subtype. Our “trihead” nanoparticle immunogen platform thus enables new insights into anti-HA immunity, establishes antigen spacing as an important parameter in structure-based vaccine design, and embodies several design features that could be used to generate next-generation vaccines against influenza and other viruses.

**HIGHLIGHTS:**

- Computational design of a closed trimeric HA head (“trihead”) antigen platform.
- Design of a rigid, extendable linker between displayed antigen and underlying protein nanoparticle enables precise variation of antigen spacing.
- Decreased antigen spacing of triheads elicits antibodies with the highest HAI, neutralizing activity, and cross-reactivity.
- Changes to antigen spacing alter epitope specificities of vaccine-elicited antibodies.

## INTRODUCTION

Influenza vaccines are of significant biomedical importance, both for the prevention of disease and as tools for studying vaccinology. The viral hemagglutinin (HA) is the key antigen of current influenza vaccines and is a well-studied model antigen. HA is a homotrimeric class I fusion protein responsible for both binding to sialoside receptors on host cells and mediating membrane fusion to deliver the viral RNA (Harrison 2015; Wu and Wilson 2020). HA consists of two distinct structural domains—the head and the stem—which have distinct functional and immunogenic properties. The head domain, comprising an upper receptor binding domain (RBD) and a lower vestigial esterase (VE) subdomain, is immunodominant, hypervariable, and the primary target of antibodies elicited by current commercial influenza vaccines (Krammer and Palese 2015; Dugan et al. 2020; Angeletti and Yewdell 2018; Guthmiller, Utset, and Wilson 2021). In contrast, the immuno-subdominant stem domain contains the membrane fusion machinery and bears broadly conserved epitopes for antibodies (Wu and Wilson 2020). Antibody responses against the HA head often potently neutralize virions by blocking receptor binding, an effect that can be measured using hemagglutination inhibition (HAI), a long-established correlate of protection (Rimmelzwaan and McElhaney 2008). However, the immunodominance of highly variable head epitopes and biased immune memory induced by early encounters with influenza antigens lead to narrow antibody specificities (Knight et al. 2020a), forcing regular updates to vaccine formulations to keep up with mutations in circulating viruses (Knight et al. 2020b; Belongia et al. 2016). Improving the potency of antibody responses could lead to direct increases in vaccine efficacy (Ortiz et al. 2023; Dunning et al. 2016).

The HA head has a variety of epitopes that have been extensively studied, revealing distinct protective mechanisms and differences in epitope conservation. Early studies with murine monoclonal antibodies (mAbs) divided the HA head into multiple antigenic sites and showed that sites nearer the sialic acid receptor binding site (RBS) mutate more frequently (Gerhard et al. 1981; Caton et al. 1982; Angeletti et al. 2017; Wiley, Wilson, and Skehel 1981). More recent studies have identified epitopes in the HA head with higher conservation that could be targeted by broadly cross-reactive vaccines. RBS-directed antibodies can broadly neutralize viruses within the same HA subtype (Bajic and Harrison 2021; Zost et al. 2019; Xu et al. 2013; Hong et al. 2013; Schmidt et al. 2013; Whittle et al. 2011)—and in some cases sporadically across subtypes (Ekiert et al. 2012; P. S. Lee et al. 2012, 2014; Li et al. 2022)—but may require avidity to achieve potency and can be sensitive to escape mutations (Zost et al. 2019). More recently, interface-directed mAbs recognizing cryptic epitopes in the interface between head domains were shown to be capable of providing broad protection across diverse influenza A subtypes in preclinical studies (Watanabe et al. 2019; Zost et al. 2021; Bangaru et al. 2019; J. Lee et al. 2016). Because these interface-directed mAbs lack neutralizing activity, effector functions are thought to play a key role in their protective capacity. Other subtype-specific epitopes throughout the head have the potential to elicit antibodies with HAI or neutralizing activity, or effector function-driven immunity (Zhu et al. 2013; Dreyfus et al. 2012; Krause et al. 2010; Li et al. 2022; Kanekiyo et al. 2019; Raymond et al. 2018; Bangaru et al. 2019; Turner et al. 2019). The location and angle of approach in part determine the protective mechanisms of head-directed antibodies, with HAI (i.e., receptor blocking) achieved by antibodies targeting apical epitopes or competing directly with binding to sialosides, while neutralizing head-directed antibodies without HAI activity often target epitopes more distant from the RBS. The diversity of epitopes in the HA head and the antibody responses they elicit offer multiple opportunities for engineering the quality of immune responses induced by next-generation vaccines.

Efforts to enhance head-directed antibody responses against specific epitopes have focused on direct modifications to epitopes in the HA head or the way it is presented to the immune system. One approach is to incorporate a diverse collection of epitopes from various viruses in single HA antigens to increase protective breadth through direct inclusion of broadly representative epitopes (Allen, Ray, and Ross 2018; Giles and Ross 2011; Sautto, Kirchenbaum, and Ross 2018). Hyperglycosylation and engineered inter-protomer disulfide bonds have been used to redirect antibody responses away from less desirable epitopes and focus them on (Bajic et al. 2019) or away from (Thornlow et al. 2021) the trimer interface. Heterologous prime-boost regimens using chimeric antigens that maintain target epitopes (Bajic et al. 2020; Caradonna et al. 2022), or alternatively diversify off-target epitopes (Sun et al. 2019; Broecker et al. 2019) or entire domains (Nachbagauer et al. 2021; Krammer et al. 2013) have also been evaluated as a mechanism for focusing antibody responses on target epitopes. Interestingly, early clinical studies with chimeric HA antigens featuring exotic head domains demonstrated the elicitation of interface-directed antibodies, perhaps because these antigens adopt open conformations that expose this epitope (Nachbagauer et al. 2021; Zhu et al. 2022). Innovative approaches for displaying HA antigens on nanoparticles have also been used to shape polyclonal antibody responses. For example, co-display of multiple HA RBD antigens of H1 subtype on ferritin nanoparticles improved antibody responses against broadly conserved epitopes within H1 HAs (Kanekiyo et al. 2019). Identifying additional strategies to maximize and shape antibody responses against target epitopes on the HA head could contribute to the development of improved influenza vaccines and inspire vaccine design for other pathogens.

The immunogenicity of a given antigen has long been linked to its spatial organization (Irvine and Read 2020). Antigen repetition and the spacing between antigens on virions have been argued to play key roles in antibody responses to infection, explaining why the immune system has evolved to recognize repetition as a danger signal (Harris et al. 2013; Klein and Bjorkman 2010; Amitai et al. 2020; Schiller and Chackerian 2014). Early studies demonstrating the importance of antigen oligomerization in the context of vaccination suggested that repetitive display every 5-15 nm may be ideally immunogenic (M. F. Bachmann and Zinkernagel 1996; M. F. Bachmann et al. 1997; Martin F. Bachmann and Jennings 2010; Dintzis, Dintzis, and Vogelstein 1976; Chackerian et al. 2002). More recent studies have used DNA origami to precisely probe the effects of antigen spacing and valency on antibody avidity and B cell activation *in vitro* (Shaw et al. 2019; Veneziano et al. 2020). Likewise, engineered series of protein nanoparticle immunogens have shown that increasing antigen valency leads to enhanced B cell activation and antibody responses *in vivo* (Kato et al. 2020; Marcandalli et al. 2019). However, defining structural correlates of immunogenicity and applying this knowledge to improve vaccine design has been challenging due to the lack of tools capable of generating systematic series of immunogens that vary specific structural features with high precision.

Here we describe a protein nanoparticle immunogen platform in which the oligomeric state and the spatial organization of displayed HA head domains can be precisely controlled. We find that both features contribute to the immunogenicity of various epitopes to shape the overall magnitude and quality of the vaccine-elicited antibody response. Our results suggest new approaches to vaccine development for influenza and other pathogens through precisely tailored nanoparticle immunogen design.

## RESULTS

### Design and Characterization of a Trihead Nanoparticle Immunogen

HA naturally forms a trimer with the stem domain containing the majority of inter-protomeric contacts, while the RBD region in the head domain has relatively fewer contacts, allowing for some flexibility or "breathing" (Benton et al. 2020; Das et al. 2018). When expressed without the stem domain, HA RBDs are monomeric when fused to other multimeric structures, such as protein nanoparticles (Kanekiyo et al. 2019). We sought to develop structure-based strategies for controlling two aspects of HA head antigen display on protein nanoparticles. First, we aimed to conformationally fix HA head antigens in either “closed” trimeric or “open” monomeric conformations, respectively hiding or exposing epitopes at the interface between head domains to determine whether each conformation has distinct immunogenic properties. Second, we aimed to study in detail the impact of antigen organization on immunogenicity by designing nanoparticle immunogens that display HA head antigens with precisely defined antigen spacing and rigidity.

As an initial step towards both goals, we aimed to use a trimeric helical bundle as a rigid, extensible linker between the displayed HA RBD antigen and an underlying protein nanoparticle scaffold. Symmetrical helical bundles can be modularly constructed from motifs such as heptad repeats to form rigid units of arbitrary length (Huang et al. 2014; Lupas, Bassler, and Dunin-Horkawicz 2017; Woolfson 2005). A mutated homotrimeric GCN4-derived helical bundle with C3 symmetry built from such repeats (PDB ID: 1GCM) was previously observed to closely match the backbone geometry of the apical portion of the HA stem that natively contacts the RBDs (Harbury, Kim, and Alber 1994; Impagliazzo et al. 2015). Furthermore, the C-terminal end of this GCN4 trimer closely matches the N-terminal helices of I53_dn5B, the trimeric component of the computationally designed icosahedral nanoparticle I53_dn5 (Ueda et al. 2020). We reasoned that using this helical bundle as a rigid connection between the RBD and I53_dn5B might stabilize the RBDs in a native-like closed trimeric state while also allowing for precise control of antigen spacing by simply modifying the number of heptad repeats.

A preliminary design was constructed by fusing the RBD (residues 56-264) of the H1N1 isolate A/New Caledonia/20/1999 (NC99) to two heptad repeats of the modified trimeric GCN4 bundle and I53_dn5B (**Figure 1A,B**). The N-terminal portion of GCN4 was redesigned to match residues in the HA stem that contact the HA head to determine if these interactions and GCN4-mediated trimerization was sufficient to drive trimeric closure of the heads. The C-terminal end of GCN4 was blended into a single continuous helix with the N terminus of I53_dn5B (referred to as the “heptad domain”) by optimally aligning the hydrophobic residues that form the core of the three-helix bundle with those of I53_dn5B. The resultant protein (referred to as “monohead”) was successfully secreted from HEK293F cells, purified by immobilized metal affinity chromatography (IMAC), and confirmed as a trimer by size exclusion chromatography (SEC) (**Figure S1A**). Biolayer interferometry (BLI) showed that the monohead component bound well to the RBS-directed mAb C05 (Ekiert et al. 2012) compared to a monomeric NC99-RBD benchmark antigen, suggesting native-like HA head folding (**Figure 1C**). However, it also bound the non-neutralizing trimer interface-directed antibody FluA-20 (Bangaru et al. 2019), which demonstrated that the interface region remained exposed and accessible for antibody recognition.

**Figure 1.**
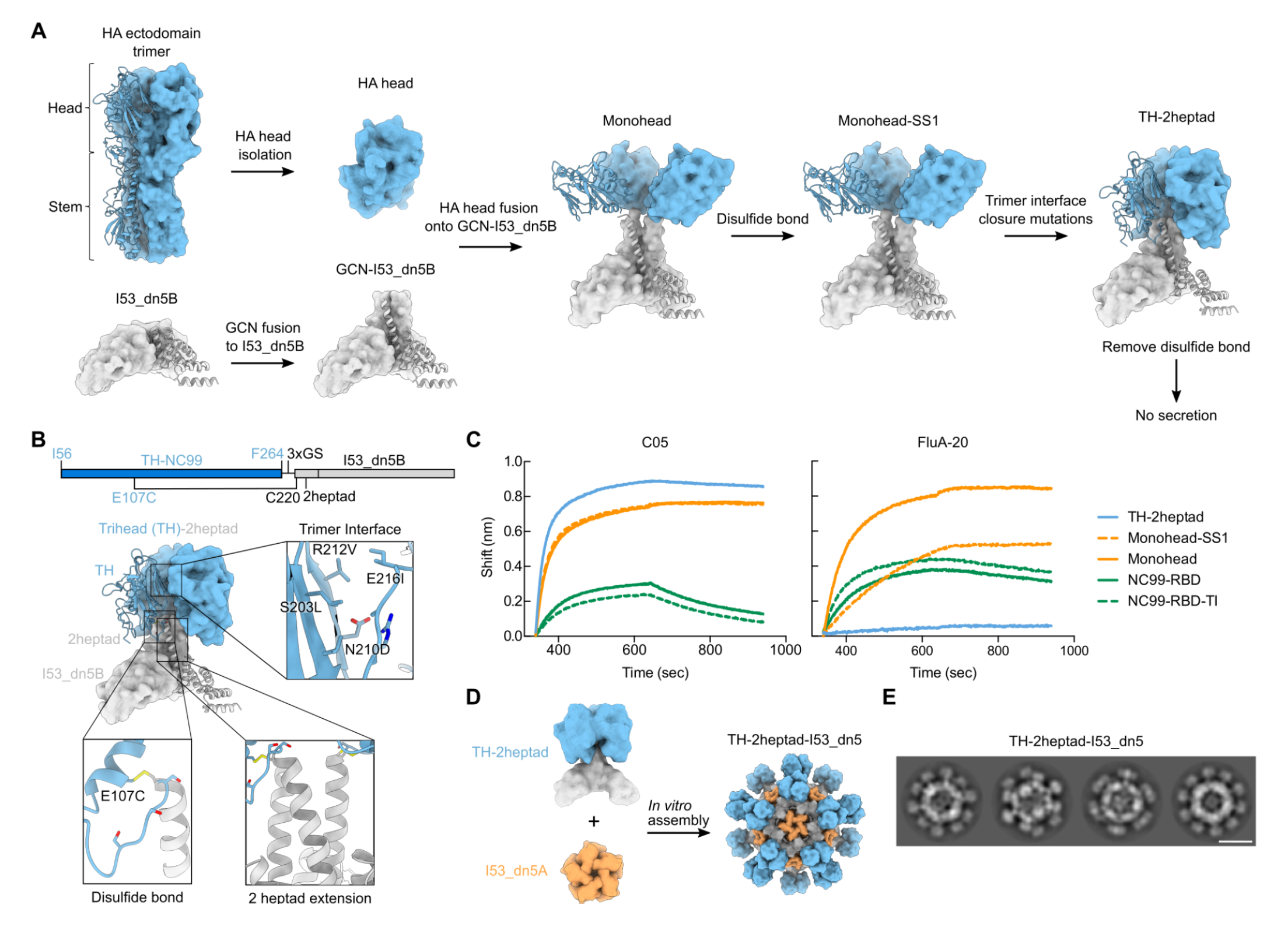
**Design and Characterization of a Trihead Nanoparticle Immunogen** (A) Schematic of design process to make TH-2heptad construct. HA-derived segments are colored in blue while segments derived from I53_dn5B and/or GCN4 are colored in gray. (B) Gene diagram and model structure of TH-2heptad with close-ups of the designed trimer interface, disulfide bond, and 2 heptad extension domain. NC99 HA numbering is in blue and trihead model numbering is in black. (C) BLI of various NC99 RBD-based constructs against C05 and FluA-20. (D) Schematic of *in vitro* assembly of the TH-2heptad-I53_dn5 nanoparticle. (E) Cryo-EM 2D class averages of TH-2heptad-I53_dn5. Scale bar = 25 nm.

We therefore designed two additional structural elements to generate closed, rigidly displayed trimeric HA heads (referred to as “triheads” (TH)). First, we used Rosetta (Leman et al. 2020) to identify a favorable intra-protomeric disulfide bond between the RBD and the N-terminal portion of the GCN4 bundle (**Figure 1A,B**). This monohead-SS1 construct was also confirmed to be a trimer by SEC but still bound to FluA-20 (**Figure 1C**). To further promote the closed trimeric state, we used Rosetta to optimize the hydrophobic interfaces between the RBDs while avoiding direct mutation of residues contacted by trimer interface-directed antibodies (Bangaru et al. 2019; Watanabe et al. 2019) (**Figure 1A,B**). A design containing the designed disulfide and four mutations, which we named TH-2heptad (S203L/N210D/R212V/E216I; **Figure 1B**), was successfully scaled up and purified by SEC, which confirmed its overall trimeric state (**Figure S1A**). In contrast to previous constructs, TH-2heptad did not bind to FluA-20, while a monomeric RBD containing the same trimer interface mutations, NC99-RBD-TI, bound as well as the wild-type NC99-RBD (**Figure 1C**). C05 binding was unaffected by the trimeric interface mutations, as expected. These results demonstrate that the trimer interface mutations help to stabilize the RBDs in a closed trimeric state, preventing FluA-20 access, but do not inherently disrupt the FluA-20 epitope.

We assembled trihead nanoparticles by mixing purified TH-2heptad trimer with the complementary pentameric component, I53_dn5A (**Figure 1D**; (Ueda et al. 2020)), and purified them using SEC (**Figures S1B,C**). Cryo-electron microscopy (cryo-EM) 2D class averages of the nanoparticles revealed clear density for trimeric antigens on the surface of the nanoparticle (**Figure 1E**). The clarity with which the triheads were resolved relative to the icosahedral nanoparticle scaffold sharply contrasts with previously described nanoparticle immunogens that used flexible genetic linkers (Marcandalli et al. 2019; Boyoglu-Barnum et al. 2021; Antanasijevic et al. 2020; Sliepen et al. 2022; Malhi et al. 2022; Walls et al. 2020; Brouwer et al. 2022) and indicates rigid attachment to I53_dn5. Removal of the disulfide between E107C and C220 in the heptad domain abolished secretion, establishing that the combination of this disulfide and the trimer interface mutations were required for trihead formation. Taken together, our BLI and cryo-EM data establish that TH-2heptad-I53_dn5 rigidly displays closed trimeric HA heads as intended.

### Design and Characterization of Trihead Nanoparticles with Varied Antigen Spacing

To enable investigation of the effects of antigen spacing on immunogenicity, we designed two new sequences that modified solely the heptad domain while retaining its rigidity. These were TH-1heptad, with one fewer heptad domain than TH-2heptad, and TH-6heptad, with four additional heptad domains that substantially lengthen the extendable region (**Figure 2A**). We also designed a fourth “bobblehead” construct that introduced a glycine-and serine-based linker between the 2heptad domain and I53_dn5B to introduce flexibility between the antigens and I53_dn5 while maintaining trimeric closure of the RBDs. Finally, we included a control component, HA-I53_dn5B, which features the full NC99 HA ectodomain fused to I53_dn5B in the same manner as we previously reported (Boyoglu-Barnum et al. 2021). Each of these novel nanoparticle components were secreted from HEK293F cells and purified via IMAC and SEC (**Figure S1A**). All of the proteins, including the original monohead component, showed similar binding to the RBS-directed mAb C05 (Ekiert et al. 2012) and the lateral patch-targeting mAb Ab6649 (Raymond et al. 2018), demonstrating the preservation of these epitopes across all designs (**Figure 2B**). By contrast, FluA-20 binding was high only for the monohead, with very slight binding to TH-6heptad and HA-I53_dn5B and no binding to any of the other constructs, indicating stable head closure. As described above for TH-2heptad, the novel components were assembled *in vitro* with the I53_dn5A pentamer to form nanoparticles and purified via SEC (**Figure S1B**). The SEC chromatograms, SDS-PAGE (**Figure S1C**), dynamic light scattering (DLS; **Figure 2C**), and negative stain electron microscopy (nsEM; **Figure 2D**) all indicated that the nanoparticles formed the intended assemblies with high homogeneity. The expected differences in size and antigen spacing between the three trihead nanoparticles were evident in the DLS and nsEM data, respectively. The lack of significant amounts of residual, unassembled component during SEC indicated efficient assembly for all trihead nanoparticles, bobblehead-I53_dn5, and HA-I53_dn5, whereas the monohead nanoparticle assembled less efficiently (**Figure S1B**; peak at ∼16 mL).

**Figure 2.**
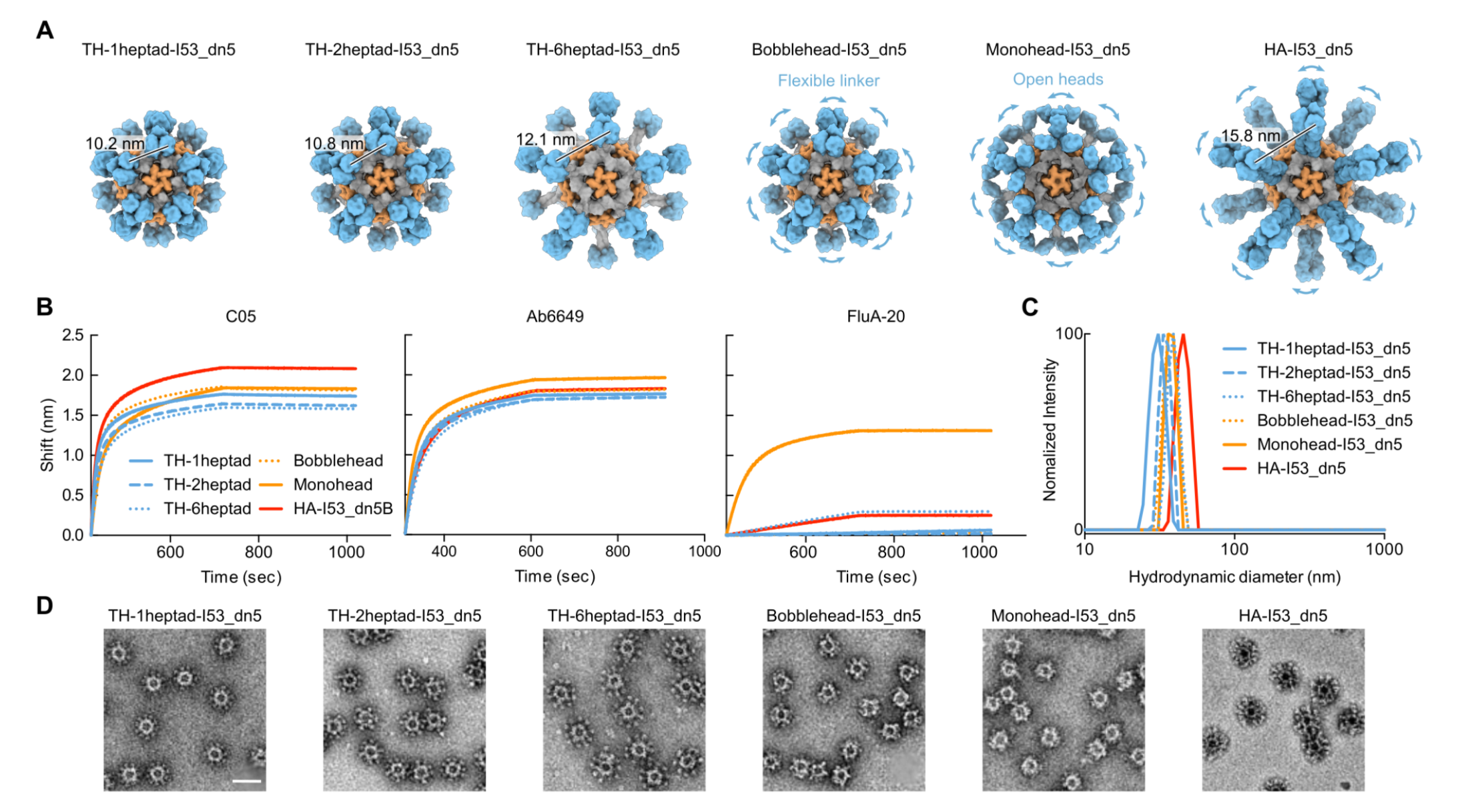
**Design and Characterization of Trihead Nanoparticles with Varied Antigen Spacing** (A) Models of trihead nanoparticles with various extension domains, monohead-I53_dn5, and HA-I53_dn5. The models are colored blue, gray and orange respectively for HA-derived segments, I53_dn5B/GCN4, and I53_dn5A. (B) BLI of trihead nanoparticle extension series components against C05, FluA-20, and Ab6649. (C) DLS of trihead nanoparticle extension series. (D) nsEM micrographs of trihead nanoparticle extension series. Scale bar = 50 nm.

### Cryo-EM of Trihead Nanoparticle Extension Series

The structural differences among the trihead, bobblehead, and monohead nanoparticles were examined using 3D reconstructions from either cryo-EM or nsEM. Cryo-EM structures were obtained for TH-1heptad-I53_dn5, TH-2heptad-I53_dn5, and TH-6heptad-I53_dn5 at respective resolutions of 4.1 Å, 6.8 Å, and 4.0 Å (**Figures 3A, S2A and S2B, Table S2**). Compared to previous cryo-EM structures of trimeric antigen-nanoparticle fusions (Boyoglu-Barnum et al. 2021; Antanasijevic et al. 2020; Marcandalli et al. 2019), the antigen density in this study was much more clearly defined, displaying the expected shape for three RBDs in the closed trimeric conformation symmetrically matched to the 3-fold symmetry axes of I53_dn5. The density observed between the RBDs and I53_dn5 was consistent with the heptad domain of each nanoparticle, with distinct lengths observed. However, the local resolution of the trihead antigens was generally lower than that of I53_dn5, indicating some degree of flexibility in the connection between the antigens and nanoparticle scaffold (**Figure S2C**). In contrast, 3D nsEM reconstructions for the bobblehead and monohead nanoparticles did not show clear density for the displayed antigens (**Figures 3A and S2D**), despite the presence of the stabilizing trimer interface mutations in bobblehead that eliminated binding to FluA-20 (**Figure 2B**). Consistent with their design, the monohead nanoparticle reconstruction revealed density related to its heptad domain rigidly attached to I53_dn5, while the reconstruction of the bobblehead nanoparticle, which contained an additional flexible linker between its heptad domain and I53_dn5, did not show such density.

**Figure 3.**
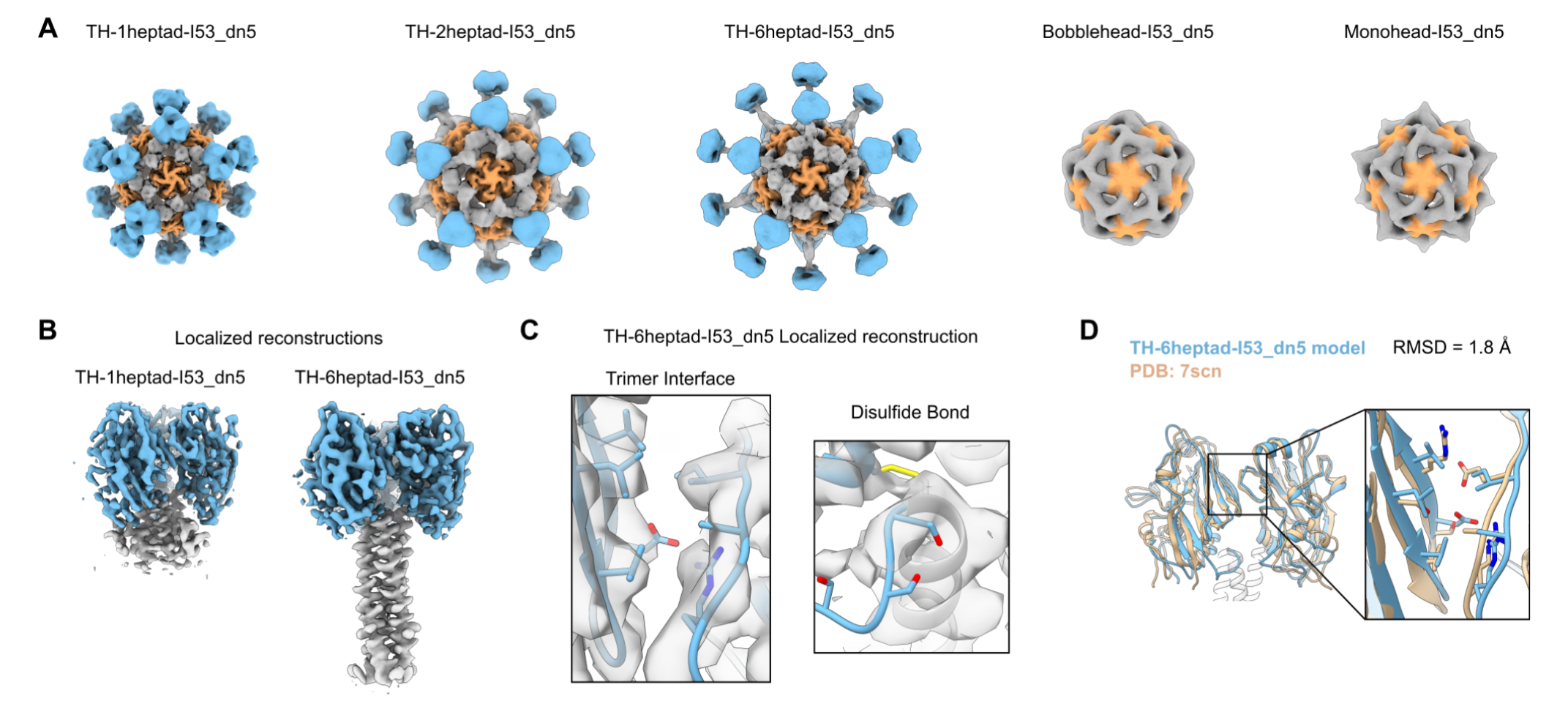
**Cryo-EM of Trihead Nanoparticle Extension Series** (A) Cryo-EM reconstructions of trihead nanoparticles and nsEM reconstructions of bobblehead and monohead nanoparticles. The densities are colored blue, gray and orange respectively for HA-derived segments, I53_dn5B/GCN4, and I53_dn5A. (B) Localized reconstructions of 1heptad and 6heptad triheads as displayed on I53_dn5 nanoparticles. (C) Close-ups of density with the corresponding built model from the TH-6heptad-I53_dn5 localized reconstruction for the engineered trimer interface and disulfide bond. Side chains for D210, R220, S266, S268, and S270 are only approximately displayed and are truncated in deposited maps. (D) Superimposition of the TH-6heptad-I53_dn5 cryo-EM model with the NC99 HA crystal structure, with a blow-up of the trimer interface where key interacting residues are displayed as sticks.

To gain a deeper understanding of the structures of the trihead antigens, localized reconstructions of the RBD and heptad domains of TH-1heptad-I53_dn5 and TH-6heptad-I53_dn5 were generated with resolutions of 3.7 Å and 3.9 Å, respectively (**Figures 3B and S2E**). These reconstructions indicated rigid connections between the RBDs and the heptad domains, suggesting that the minor flexibility observed between the RBDs and the underlying nanoparticle likely derives from the heptad domains and their connections to I53_dn5, or is spread out across these domains. The atomic models of TH-1heptad and TH-6heptad antigens generated from these localized reconstructions confirmed the desired structural effects from the designed stabilizing features. Specifically, the inter-RBD interface showed hydrophobic packing between the designed mutations, and density consistent with disulfide formation was observed between E107C and GCN4, with less clear density where the flexible genetic linker was expected between each RBD and heptad domain (**Figures 3C**). The backbone atoms from all three TH-6heptad RBDs were structurally aligned with the RBDs from a cryo-EM structure of the NC99 trimeric ectodomain (PDB ID: 7SCN), revealing close similarity between the structures (1.8 Å Cα RMSD; **Figure 1D**). These detailed structural analyses complement our biophysical and antigenic data to confirm that the nanoparticle immunogens exhibit both of the target structural characteristics: stabilization of the RBD antigen in a native-like closed trimeric state and a rigid connection to the I53_dn5 nanoparticle that allows for variable antigen spacing.

### Antibody Responses in Mice Immunized with Trihead Nanoparticle Extension Series

We next evaluated the immunogenicity of our trihead nanoparticle extension series in mice. In a first study, groups of 10 BALB/c mice were immunized with AddaVax-adjuvanted protein at weeks 0, 4, and 8 (**Figure S3A**). Vaccine-matched NC99 HA binding titers in week 10 sera were comparably high across all groups, with nearly all samples surpassing the upper limit of detection of the assay (**Figure S3B**). However, some differences in microneutralization titers were observed, with TH-1heptad-I53_dn5 and HA-I53_dn5 eliciting equivalent levels of neutralizing activity that were significantly higher than TH-6heptad-I53_dn5 and bobblehead-I53_dn5 (**Figure S3C**). Neutralizing activity within the trihead extension series trended toward decreasing with increasing heptad region length. To expand the dynamic range of our serological readouts, we performed a similar study using unadjuvanted immunogens (**Figure 4A**). Analyzing immune sera from week 10 showed that TH-1heptad-I53_dn5 elicited significantly higher NC99 HAI titers than all other groups, including 5.3-fold higher than TH-6heptad-I53_dn5 and 5.2-fold higher than HA-I53_dn5 (**Figure 4B**). The pattern of microneutralization titers across the groups in this study trended similarly to the adjuvanted study, with the TH-1heptad-I53_dn5 and HA-I53_dn5 sera again having the highest neutralizing activity (**Figure 4C**). Lastly, in a second adjuvanted study we analyzed binding titers in week 10 sera against three vaccine-mismatched H1 HAs (**Figure 4A and D**). The highest cross-reactive antibody titers were induced by monohead-I53_dn5 and HA-I53_dn5, possibly due to antibodies against conserved epitopes in the trimer interface and non-RBD domains, respectively. However, only these two nanoparticle immunogens elicited higher levels of cross-reactive antibodies than TH-1heptad-I53_dn5, which in turn elicited higher vaccine-mismatched binding titers than all of the other trihead nanoparticles, though not always with statistical significance. Taken together, these results demonstrate a consistent trend toward increased serum antibody potency and breadth with decreased trihead antigen spacing, with the TH-1heptad nanoparticle eliciting significantly more potent antibody responses than the other trihead and monohead immunogens. Furthermore, the high levels of neutralizing activity and cross-reactive antibodies elicited by HA-I53_dn5 clearly indicate that deletion of domains outside the RBD can eliminate epitopes that aid in eliciting neutralizing and cross-reactive antibodies. Nevertheless, the dense display of closed trimeric head domains in TH-1heptad-I53_dn5 provides an alternative approach to eliciting potent functional antibody responses, most notably through induction of higher levels of antibodies with HAI activity.

**Figure 4.**
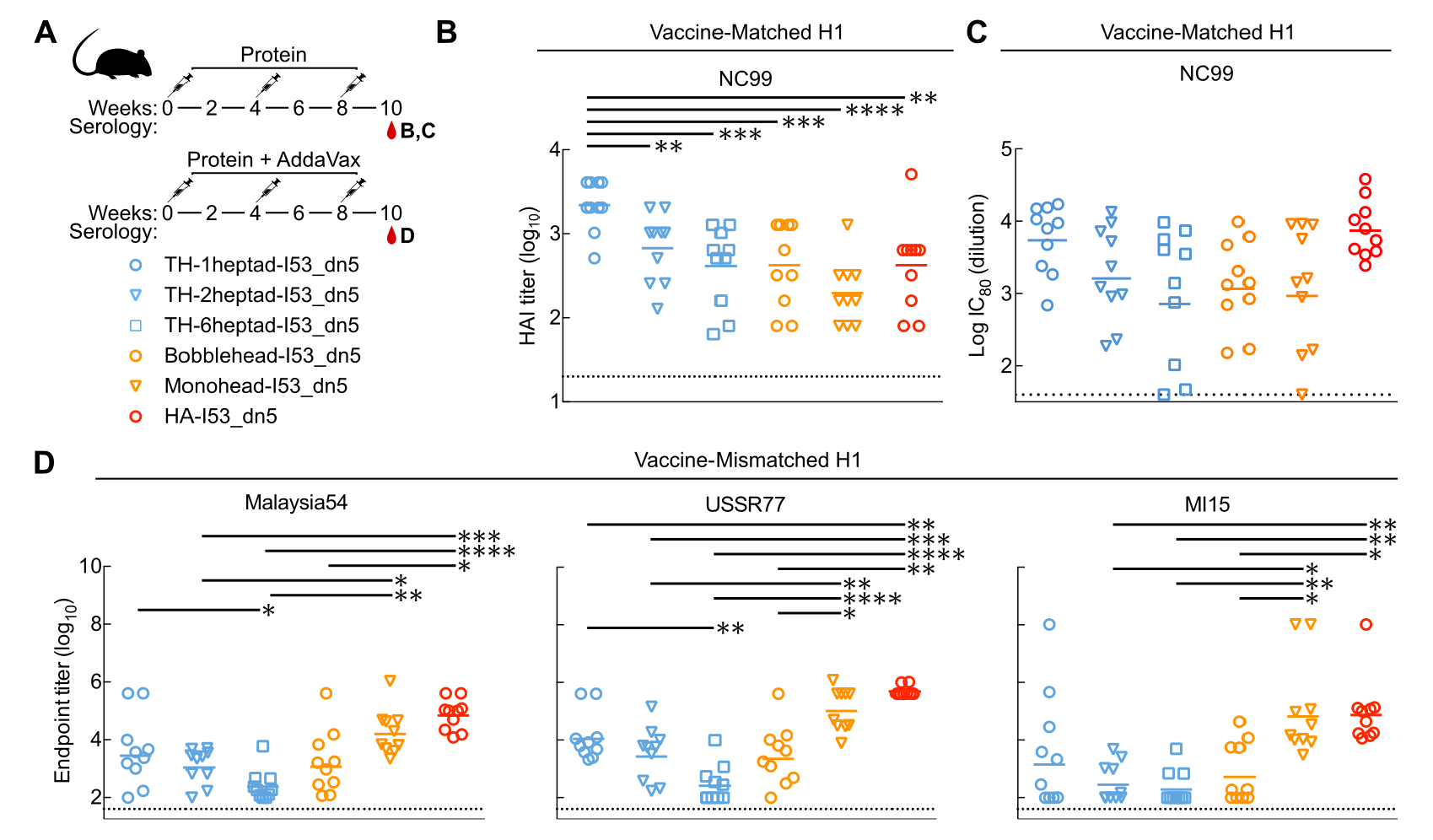
**Antibody Responses in Mice Immunized with Trihead Nanoparticle Extension Series** (A) Mouse immunization schedule and groups for trihead nanoparticle extension series that were either unadjuvanted or adjuvanted. (B) Vaccine-matched HAI titers in unadjuvanted immune sera. (C) Vaccine-matched microneutralization titers in unadjuvanted immune sera. (D) Vaccine-mismatched H1 ELISA binding titers in adjuvanted immune sera. Statistical significance was determined using one-way ANOVA with Tukey’s multiple comparisons test; ∗p < 0.05; ∗∗p < 0.01; ∗∗∗p < 0.001; ∗∗∗∗p < 0.0001.

### Epitope Mapping of Vaccine-elicited Antibodies

We generated NC99-based HA probes featuring knockout mutations in specific epitopes to map the epitope specificities of vaccine-elicited antibodies. One set of probes comprised the trimeric HA ectodomain fused to foldon (referred to as NC99), with or without epitope knockout mutations. One RBS-knockout probe was designed by introducing a single L194W mutation into the NC99 probe, replacing a highly conserved leucine residue in the center of the RBS with a bulkier tryptophan residue (**Figure 5A**). A second RBS-knockout probe introduced an N-linked glycan within the RBS via T155N and K157T mutations. We expected the single L194W mutation to be more sensitive to antibodies that bind directly within the RBS pocket, and that the bulky engineered glycan at residue 155 would interfere with binding to both RBS-directed and RBS-proximal antibodies. We also introduced the trimer interface mutations used in the trihead antigen into the NC99 ectodomain trimer to maximally stabilize HA in the closed state (NC99-CS), as HA ectodomains are thought to breathe and transiently expose epitopes at the trimer interface (Turner et al. 2019; Bangaru et al. 2019).

**Figure 5.**
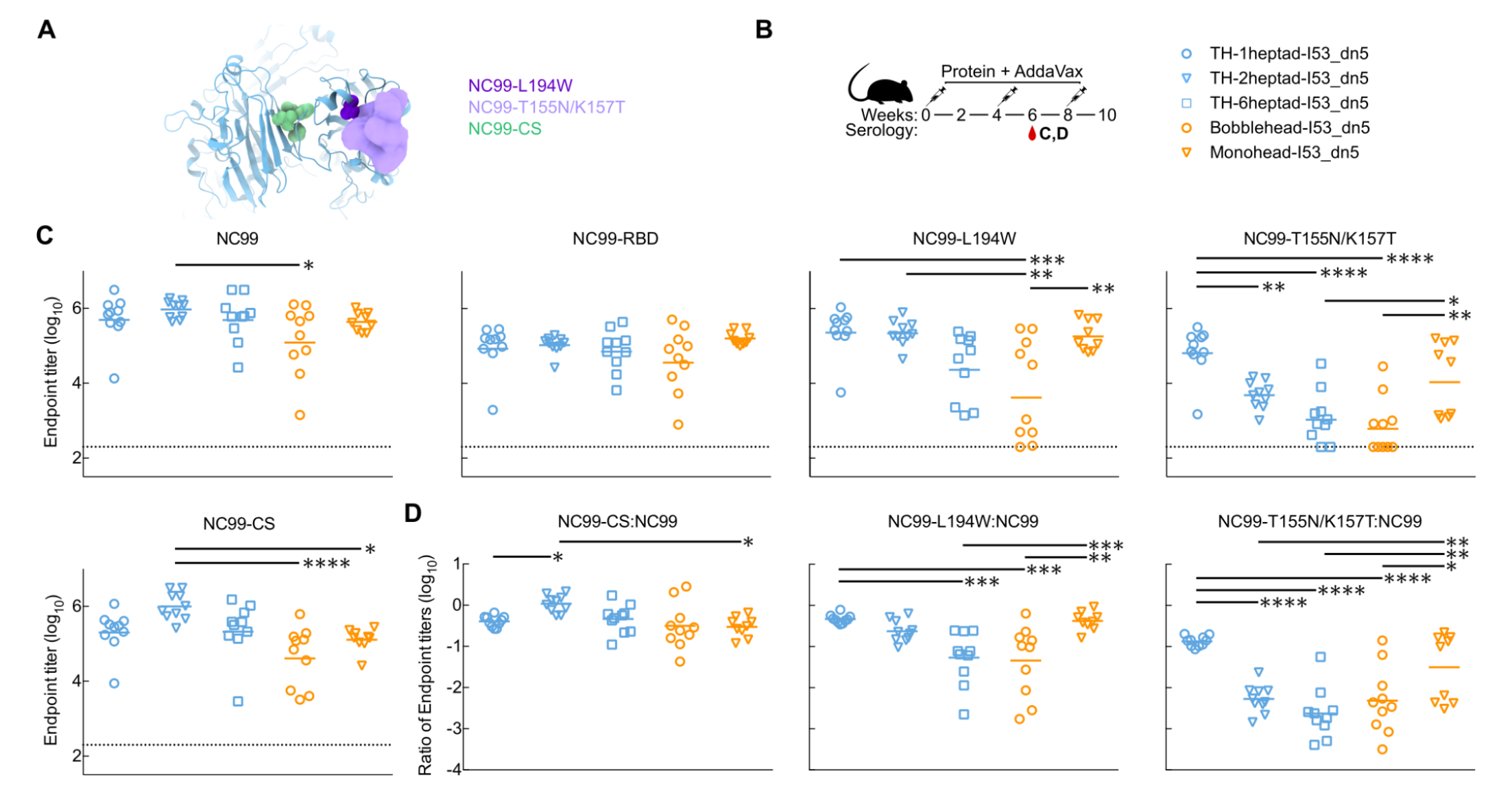
**Epitope Mapping of Vaccine-elicited Antibodies** (A) Structural schematic of NC99 probes. Sphere representations show mutated residues. For NC99-T155N/K157T, density of a modeled glycan is shown. (B) Mouse immunization schedule and groups for adjuvanted trihead nanoparticle extension series. (C) ELISA binding titers against various NC99 probes in week 6 adjuvanted immune sera. (D) Ratios of ELISA binding titers of mutated NC99 probes to NC99 in panel B. Statistical significance was determined using one-way ANOVA with Tukey’s multiple comparisons test; ∗p < 0.05; ∗∗p < 0.01; ∗∗∗p < 0.001; ∗∗∗∗p < 0.0001.

Finally, the wild-type monomeric NC99 RBD was also produced to allow detection of antibodies binding anywhere on the HA head, including the trimer interface. Antigenic characterization of these probes was first performed using BLI. All of the probes bound to the lateral patch-targeting mAb Ab6649 (Raymond et al. 2018), indicating proper folding of the head domain (**Figure S4A**). As expected, all ectodomain trimer probes also bound to the anti-stem mAb CR9114 (Dreyfus et al. 2012), whereas the NC99-RBD did not. FluA-20 robustly bound the NC99-RBD, less efficiently bound NC99 and NC99-T155N/K157T, and did not bind NC99-L194W or NC99-CS, confirming that the trimer interface epitope is accessible in NC99-RBD but not NC99-CS. C05 bound NC99, NC99-CS, and NC99-RBD, but did not bind either NC99-L194W or NC99-T155N/K157T, indicating that these modifications to the RBS prevent RBS-directed antibody binding as intended. All four trimeric ectodomain probes appeared mostly as small elongated rods by nsEM (**Figure S4B**), consistent with side views of HA ectodomain trimers. Some rosette formation was observed in all samples, especially NC99-T155N/K157T, as frequently occurs with recombinant HA trimers (Laver and Valentine 1969; McCraw, Gallagher, and Harris 2016). Taken together, these results demonstrate that each probe possesses the desired structure and antigenicity.

These NC99 probes were then used to roughly estimate the epitope specificities of serum antibodies elicited by each immunogen. We used ELISA to measure the levels of antibodies against each probe in week 6 sera from our first adjuvanted study (**Figures 5B** and **S3**). All of the immunogens elicited similarly high binding titers against NC99, with the exception of bobblehead-I53_dn5, which was slightly lower (**Figure 5C**). The same trend was observed in the NC99-RBD binding titers. Interestingly, the pattern of titers against the NC99-L194W and NC99-T155N/K157T RBS knockout probes mirrored the HAI and neutralization data (**Figures 4B-C** and **S3C**) in that the titers elicited by the trihead nanoparticle immunogens decreased with increasing heptad region length, while bobblehead-I53_dn5 elicited low and variable levels of antibodies against these probes. This pattern suggests that TH-1heptad elicited higher levels of receptor-blocking and neutralizing antibodies that bind epitopes outside the RBS (e.g., 2D1 (Xu et al. 2010; Yu et al. 2008)). Ratiometric analysis of the binding of each serum sample to the two RBS knockout probes compared to NC99 is consistent with this suggestion (**Figure 5D**). Sera from mice that received monohead-I53_dn5 also appeared relatively insensitive to the RBS knockout mutations, although a subset of the sera showed substantially less binding to NC99-T155N/K157T (**Figure 5C and D**). Interestingly, these sera did not show significantly reduced binding to the NC99-CS probe, in which the trihead interface mutations reduce exposure of the FluA-20 epitope (**Figure 5C and D**). Nevertheless, the insensitivity to the RBS knockout mutations and cross-reactive antibody binding data in **Figure 4D** suggest that antibody responses against the trimer interface are elicited by monohead-I53_dn5. This conclusion is supported by our accompanying manuscript (Dosey et al., n.d.), which shows that hyperglycosylation of monohead-I53_dn5 can further increase the magnitude of the antibody response against the trimer interface relative to other epitopes on the RBD. In all, our data support the hypothesis that antigen display geometry can shape the magnitude and epitope specificities of vaccine-elicited antibodies, with TH-1heptad_I53_dn5 eliciting potent receptor-blocking antibody responses that nevertheless target a wider variety of epitopes compared to similar immunogens that vary only with respect to antigen display geometry.

## DISCUSSION

Here we designed a native-like trimeric HA head antigen that can be rigidly displayed on a protein nanoparticle scaffold and explored the effects of antigen display geometry on vaccine-elicited antibody responses. Combining domain-based antigens with nanoparticle display is an established and powerful strategy for enhancing the potency and breadth of antibody responses against epitopes that are targeted by potently neutralizing or protective antibodies. Domain-based antigens are small and often monomeric, making nanoparticle display particularly advantageous: in addition to improving their interactions with B cells (Veneziano et al. 2020; Shaw et al. 2019; Abbott et al. 2018; Kato et al. 2020; Martin F. Bachmann and Jennings 2010), it also enhances their trafficking and localization *in vivo* by virtue of the increased size of the immunogen and in some cases dense display of N-linked glycans (Tokatlian et al. 2019; Martin et al. 2020; Read et al. 2022). The SARS-CoV-2 Spike RBD is a prominent example of a small monomeric antigen whose immunogenicity is greatly enhanced by multivalent display on protein nanoparticles (Walls et al. 2020; Cohen et al. 2021; Dalvie et al. 2021), and the recent licensure of an RBD nanoparticle vaccine for SARS-CoV-2 has clinically validated this approach (Song et al. 2022, 2023). Additional examples of nanoparticle display of domain-based antigens focusing responses on key neutralizing epitopes include Epstein-Barr virus gp350 (Kanekiyo et al. 2015) and human cytomegalovirus glycoprotein B (Perotti et al. 2020). The strategy has also been combined with antigen stabilization (Ellis et al. 2021; McLeod et al. 2022; Dalvie et al. 2021), germline targeting (Jardine et al. 2013; Duan et al. 2018; Leggat et al. 2022), and mosaic nanoparticle display (Cohen et al. 2021, 2022; Walls et al. 2021; Kanekiyo et al. 2019) to further increase the potency and breadth of vaccine-elicited antibodies.

Displaying the influenza HA head in its native-like trimeric conformation required substantial structure-based antigen design. Our strategy was similar to those used to stabilize HA stem and coronavirus Spike S2 antigens by deleting undesired domains and then repairing the resulting conformational changes or instabilities with stabilizing mutations or topological rearrangements (Yassine et al. 2015; Corbett et al. 2019; Impagliazzo et al. 2015; Hsieh et al. 2021; Bowen et al. 2022). We focused on restoring the native-like closed trimeric arrangement of the displayed RBDs by strengthening the interactions between protomers, an approach that has been used to stabilize other oligomeric antigens in native-like conformations (Impagliazzo et al. 2015; Ellis et al. 2022; Milder et al. 2022; Joyce et al. 2016; Che et al. 2023; Stewart-Jones et al. 2021; Hsieh et al. 2022; Chuang et al. 2017; de Taeye et al. 2015; Rutten et al. 2018). Trimer closure required both computationally designed hydrophobic interactions at the trimer interface and a rigid, disulfide-mediated connection to an underlying trimerization domain. In our accompanying manuscript we demonstrate that this strategy can be generalized to multiple H1 HAs and combined with additional antigen engineering to further refine vaccine-elicited antibody responses (Dosey et al., n.d.). These two features combine to establish triheads as a robust and versatile antigen platform for influenza vaccine design. In the future, similar design approaches may be useful for stabilizing other oligomeric domain-based antigens in native-like conformations.

In contrast to previous studies that aimed to eliminate immunodominant domains to focus responses on conserved subdominant epitopes (Impagliazzo et al. 2015; Yassine et al. 2015; Corbett et al. 2019), our goal was to investigate how altering display geometry could influence antibody responses to a complex, domain-based antigen. To achieve this, we needed a system where display geometry could be modulated without affecting other vaccine characteristics that could impact antibody responses. Such a system requires a connection to the underlying nanoparticle that is both rigid and extendable. Previous work showed that DARPin proteins can be used as rigid adaptors between protein nanostructures and smaller monomeric proteins to facilitate structure determination by cryo-EM, but did not allow for precise alteration of display geometry (Liu, Huynh, and Yeates 2019; Vulovic et al. 2021). We developed a nanoparticle component featuring an extendable N-terminal three-helix coiled-coil and designed a rigid attachment between the trihead antigen and the coiled-coil that, to our knowledge, provides the most rigid and tunable connection between an antigen and protein nanoparticle scaffold to date. Although DNA origami has been used to precisely probe the effects of antigen organization on B cell activation *in vitro* (Shaw et al. 2019; Veneziano et al. 2020), the instability of DNA *in vivo* has limited evaluation of such structures in a fully functioning immune system. Elegant work displaying engineered HIV outer domain antigens on a series of protein nanoparticle scaffolds with different symmetries and valencies showed that responses to vaccination depend strongly on these structural factors and are driven by interactions with B cells (Kato et al. 2020). Our findings that altering antigen spacing alone can affect the HAI, microneutralization, cross-reactivity, and epitope specificity of vaccine-elicited antibodies further support that antigen organization is an important determinant of immunogenicity and establish that it can be optimized through structure-based antigen design.

There are several limitations to our study that present opportunities for future research. First, our serological data do not allow us to identify the mechanistic basis for the effects of antigen spacing on B cell recognition and antibody responses. Additionally, the icosahedral symmetry of I53_dn5 results in various geometric relationships between the displayed triheads (e.g., between neighboring antigens across the icosahedral two-fold or five-fold symmetry axes), and it is unclear which relationships are most important for interactions with antigen-specific B cells. Furthermore, additional studies will be required to determine whether structural correlates of immunogenicity generalize across antigens since each antigen has unique structural properties. Finally, there are many factors beyond direct immunogen-B cell interactions that could affect the immunogenicity of our nanoparticle immunogen series *in vivo*, including antigen trafficking (Tokatlian et al. 2019; Martin et al. 2020; Read et al. 2022), conformational stability (Pauthner et al. 2017; Cirelli et al. 2019), and physical integrity (Aung et al. 2023), among others. To address these limitations, future studies will need to evaluate additional series of immunogens that systematically alter specific immunogen properties, an endeavor that will be facilitated by powerful new machine learning methods for protein structure prediction and design (Dauparas et al. 2022; Watson et al. 2022; Baek et al. 2021; Lutz et al. 2023). Corresponding advances in animal models of vaccination are providing increasingly precise measurements of immunogen performance *in vivo* (Kratochvil et al. 2021; Melzi et al. 2022; Luo et al. 2023; X. Wang et al. 2021; Kato et al. 2020). Combining these technologies to define structural correlates of immunogenicity could be a powerful route to improved vaccines against influenza and many other pathogens.

## METHODS

### Structure-based Design

All design and modeling was performed using Rosetta and PyMOL. Briefly, modeling of linker regions was performed using RosettaRemodel (Huang et al. 2011). Disulfide design between the HA head and the GCN4-based rigid linker region (PDB 1GCM) was assessed using the Disulfidize mover after symmetrically superimposing HA head domains (PDB 5UG0) on the rigid linker region based on the native orientation of HA heads to the apical region of the HA2 domain, which is structurally similar to GCN4. To determine stabilizing mutations at the HA head interface, Rosetta design was performed on residues 203, 210, 212 and 216 of PDB 5UG0 using resfile-based design methods described previously (Ellis et al. 2021, 2022) with C3-symmetry imposed to model the closed state of the HA heads. Mutations were then transferred to the NC99 sequence to complete the design sequence.

### Gene Expression and Protein Purification

All HA constructs used in this study contained the Y98F mutation to reduce aggregation (Whittle et al. 2014), were codon-optimized for human cell expression, and cloned into the CMV/R vector (Barouch et al. 2005) by Genscript with a C-terminal hexahistidine affinity tag. PEI MAX was used for transient transfection of HEK293F cells. After four days, mammalian cell supernatants were clarified via centrifugation and filtration. Monohead and trihead components and HA foldons were all purified using IMAC. 1 mL of Ni^2+^-sepharose Excel or Talon resin was added per 100 mL clarified supernatant along with 5 mL of 1 M Tris, pH 8.0 and 7 mL of 5 M NaCl and left to batch bind while shaking at room temperature for 30 min. Resin was then collected in a gravity column, washed with 5 column volumes of 50 mM Tris, pH 8.0, 500 mM NaCl, 20 mM imidazole, and protein was eluted using 50 mM Tris, pH 8.0, 500 mM NaCl, 300 mM imidazole. Further component purification was done using SEC on a Superdex 200 Increase 10/300 gel filtration column equilibrated in 25 mM Tris, pH 8.0, 150 mM NaCl, 5% glycerol.

Expression and purification of the I53_dn5A pentamer component from *E. coli* was carried out as previously described (Boyoglu-Barnum et al. 2021) using the stabilized I53_dn5A.1 sequence (Ueda et al. 2020). Assembly of all nanoparticles was carried out by mixing HA-I53_dn5B-based components and pentameric I53_dn5A together *in vitro* at a 1:1 molar ratio at 15-40 μM final concentrations. Nanoparticles were left to assemble for 30 min at room temperature with rocking. Nanoparticles were then purified using SEC on a Superose 6 Increase 10/300 gel filtration column equilibrated in 25 mM Tris, pH 8.0, 150 mM NaCl, 5% glycerol.

Following purification, nanoparticle quality and concentration was first measured by UV-vis spectroscopy. Nanoparticle polydispersity and purity was then assessed using SDS-PAGE, DLS, and nsEM. Finally, endotoxin levels were measured using the LAL assay, with all immunogens used in animal studies containing less than 100 EU/mg in the final dose. Final immunogens were flash-frozen using liquid nitrogen and stored at −80°C.

### Bio-layer Interferometry (BLI)

BLI was carried out using an Octet Red 96 system, at 25°C with 1000 rpm shaking. Anti-HA antibodies were diluted in kinetics buffer (PBS with 0.5% serum bovine albumin and 0.01% Tween) to a final concentration of 10 μg/mL before loading onto protein A biosensors (Sartorius) for 200 s. Proteins were diluted to 400-500 nM in kinetics buffer and their association was measured for 200 s, followed by dissociation for 200 s in kinetics buffer alone.

### Dynamic Light Scattering

DLS was carried out on an UNcle (UNchained Labs) at 25°C. 10 acquisitions of 5 sec each were acquired for each spectrum. Protein concentration (ranging from 0.1-1 mg/mL) and buffer conditions were accounted for in the software.

### Negative stain EM

Nanoparticle samples were diluted to 0.01 mg/mL immediately prior to adsorption to glow-discharged carbon-coated copper grids for about 60 seconds prior to a 2% uranyl formate staining. Micrographs were recorded using the Leginon (Suloway et al. 2005) on a 120 KV FEI Tecnai G2 Spirit with a Gatan Ultrascan 4000 4k × 4k CCD camera at 67,000× nominal magnification. The defocus ranged from −1.0 to −2.0 µm and the pixel size was 1.6 Å.

### Cryo-EM Sample Preparation, Data Collection, and Data Processing

Three microliters of 3 mg/mL TH-1heptad-I53_dn5, TH-2heptad-I53_dn5 or TH-6heptad-I53_dn5 nanoparticle samples were loaded onto freshly glow discharged R 2/2 UltrAuFoil grids, prior to plunge freezing using a Vitrobot Mark IV (ThermoFisher Scientific) with a blot force of 0 and 6 sec blot time at 100% humidity and 22°C. For the TH-1heptad-I53_dn5 and TH-6heptad-I53_dn5 data sets, data were acquired on an FEI Glacios transmission electron microscope operated at 200 kV equipped with a Gatan K2 Summit direct detector. Automated data collection was carried out using Leginon (Suloway et al. 2005) at a nominal magnification of 36,000× with a pixel size of 1.16 Å. The dose rate was adjusted to 8 counts/pixel/s, and each movie was acquired in counting mode fractionated in 50 frames of 200 ms. 1,274 and 1,647 micrographs were collected with a defocus range between −0.5 and −2.5 μm. For the TH-2heptad-I53_dn5 data set, data were acquired using an FEI Titan Krios transmission electron microscope operated at 300 kV and equipped with a Gatan K2 direct detector and Gatan Quantum GIF energy filter, operated in zero-loss mode with a slit width of 20 eV. Automated data collection was carried out using Leginon (Suloway et al. 2005) at a nominal magnification of 130,000× with a pixel size of 0.525 Å. 224 micrographs were collected with a defocus range comprised between −0.5 and −2.5 μm, respectively. The dose rate was adjusted to 15 counts/pixel/s, and each movie was acquired in super-resolution mode fractionated in 75 frames of 40 ms. Movie frame alignment, estimation of the microscope contrast-transfer function parameters, particle picking, and extraction were carried out using Warp (Tegunov and Cramer 2019). Two rounds of reference-free 2D classification were performed using CryoSPARC (Punjani et al. 2017) to select well-defined particle images. These selected particles were subjected to two rounds of 3D classification with 50 iterations each (angular sampling 7.5° for 25 iterations and 1.8° with local search for 25 iterations) using Relion (Zivanov et al. 2018) with an initial model generated with ab-initio reconstruction in CryoSPARC (Punjani et al. 2017). 3D refinements with icosahedral symmetry were carried out using non-uniform refinement along with per-particle defocus refinement in CryoSPARC. Selected particle images were subjected to the Bayesian polishing procedure (Zivanov, Nakane, and Scheres 2019) implemented in Relion 3.1 before performing another round of non-uniform refinement in cryoSPARC followed by per-particle defocus refinement and again non-uniform refinement. To further improve the density of the trimeric HA head, a localized reconstruction method was used (Ilca et al. 2015). The designed asymmetric unit models were fitted to the icosahedral map and the locations of 20 HA head trimers in the nanoparticles were manually defined in UCSF Chimera by placing a marker in the center of each HA head trimer. A total of 1,452,820 sub-particles and 718,940 sub-particles were selected for TH-1heptad HA trihead and TH-6heptad HA trihead respectively using Scipion (Abrishami et al. 2021). The selected sub-particles were extracted in a box of 180 × 180 pixels from micrographs and subjected to one round of 3D classification using Relion (Zivanov et al. 2018). Particles belonging to classes with the best resolved trimeric HA trihead density were selected and then subjected to local refinement with C3 symmetry using CryoSPARC. Local resolution estimation and sharpening were carried out using CryoSPARC. Reported resolutions are based on the gold-standard Fourier shell correlation (FSC) of 0.143 criterion and Fourier shell correlation curves were corrected for the effects of soft masking by high-resolution noise substitution (Chen et al. 2013; Rosenthal and Henderson 2003).

### Model Building and Refinement

UCSF Chimera (Pettersen et al. 2004) and Coot (Emsley et al. 2010) were used to build atomic models that fit into the cryoEM density maps. TH-1heptad and TH-6heptad HA trihead models were refined and relaxed using Rosetta using sharpened and unsharpened maps (Frenz et al. 2019; R. Y.-R. Wang et al. 2016).

### Animal experiments

All animal experiments were reviewed and approved by the Institutional Animal Care and Use Committee of the VRC, NIAID, NIH. All animals were housed and cared for in accordance with local, state, federal, and institutional policies of NIH and American Association for Accreditation of Laboratory Animal Care. The space temperature in the rodent facility is set to 22°C ± 3 degrees. The humidity is maintained between 30% and 70%. The automatic light cycle is a 12 h on/off photo-period.

### Immunization

Female BALB/c mice (Jackson Laboratories) were immunized intramuscularly with 22 µmol purified nanoparticle immunogen in the presence or absence of AddaVax™ (InvivoGen) at weeks 0, 4, and 8. Formulated nanoparticle immunogens in 50 μl were given into each hind leg. Serum samples were collected before and after each immunization and used for immunological assays. (**Figure 4, 5 and S3**).

### ELISA

Antigen-specific IgG levels in immune sera were measured by ELISA. The plates were coated with 2 μg ml^−1^ of recombinant HA-foldon proteins and incubated at 4°C overnight. Plates were then blocked with PBS containing 5% skim milk at 37°C for 1 h. mAbs and immune sera were serially diluted in four-fold steps and added to the wells for 1 h. Horseradish peroxidase (HRP)-conjugated anti-human (SouthernBiotech, Catalog 2040-05, used 1/5,000); anti-mouse IgG (SouthernBiotech, Catalog 1080-05, used 1/5000) antibody was added and incubated at 37°C for 1 h. The wells were developed with 3,3′,5′,5-tetramethylbenzidine (TMB) substrate (KPL), and the reactions were stopped by adding 1 M H_2_SO_4_ before measuring absorbance at 450 nm with a Spectramax Paradigm plate reader (Molecular Devices). Sera from mice immunized with PBS or an irrelevant antigen (DS-Cav1-I53_dn5; (Ueda et al. 2020)) were used as negative controls, and did not yield signal above background.

### HAI

Influenza antibody titers were detected using HAI (Hemagglutination Inhibition) with 2-fold serial dilutions of RDE-treated mouse serum as described elsewhere (World Health Organization and WHO Global Influenza Surveillance Network 2011). Briefly, one part serum sample was mixed with three parts of diluted receptor destroying enzyme (RDE II; Denka Seiken) and treated at 37°C for 16-18 hours followed by heat inactivation at 56°C for 40 min. RDE-treated serum was then serially diluted in a 96-well format with PBS (total final volume of 25 μL per dilution) and 25 μL of A/New Caledonia/20/1999 R3ΔPB1 virus (at a concentration of 8HA) was added to each well with mixing. After a 1-hour incubation at room temperature, 50 μL of 0.5% washed turkey whole blood (Lampire, cat# 7209403) was added to each well. The samples were allowed to incubate an additional 30 minutes at room temperature, and the serum dilution that no longer successfully inhibited hemagglutination as seen by drip test was documented as the HAI titer for each sample.

### Reporter-based Microneutralization Assay

Influenza A Reporter viruses were made as previously described (Creanga et al. 2021). Briefly, H1N1 viruses were made with a modified PB1 segment expressing the TdKatushka reporter gene (R3ΔPB1), rescued, and propagated in MDCK-SIAT-PB1 cells in the presence of TPCK-treated trypsin (1 μg ml^−1^, Sigma) at 37 °C. Virus stocks were stored at −80 °C and were titrated before use in the assay. Mouse sera were treated with receptor destroying enzyme (RDE II; Denka Seiken) and heat-inactivated before use in neutralization assays. 384-well plates (Grenier) were pre-seeded with 1.0×10^5^ MDCK-SIAT1-PB1 cells and incubated overnight. Immune sera or monoclonal antibodies (FluA-20, Ab6649, C05) were serially diluted and incubated for 1 h at 37 °C with pre-titrated virus (A/New Caledonia/20/1999). Serum-virus mixtures were then transferred in quadruplicate onto the pre-seeded 384-well plates and incubated at 37 °C for 18-26 hours. The number of fluorescent cells in each well was counted automatically using a Celigo image cytometer (Nexcelom Biosciences). IC_80_ values, defined as the serum dilution or antibody concentration that gives 80% reduction in virus-infected cells, were calculated from neutralization curves using a four-parameter nonlinear regression model.

### Lead Contact

Further information and requests for resources and reagents should be directed to and will be fulfilled by the Lead Contact, Neil King.

## Acknowledgements

This work was funded by a generous gift from Open Philanthropy (N.P.K.); the Audacious Project at the Institute for Protein Design (N.P.K.); the Defense Threat Reduction Agency (HDTRA1-18-1-0001 to N.P.K.); the National Institute of Allergy and Infectious Diseases (P01 AI167966 to D.V. and N.P.K.); the intramural research program of the Vaccine Research Center, National Institute of Allergy and Infectious Diseases, National Institutes of Health (M.K. and B.S.G.); the University of Washington Arnold and Mabel Beckman Cryo-EM Center; and the Frederick National Laboratory for Cancer Research, NIH, under Contract HHSN261200800001 (Y.T.).

## Author Contributions

D.E., A.D., S.B., B.S.G, M.K., and N.P.K. designed experiments. D.E., A.D., and N.P.K. wrote the paper. G.U. and J.A.F. designed the I53_dn5 nanoparticle. D.E. designed triheads and gene constructs. D.E., M.M., D.P., N.M., and S.C. carried out protein production, with supervision from L.C. D.E. and A.D. carried out BLI and DLS. D.E., A.D., and Y.P. performed the nsEM. Y.P. and Y.T. performed cryoEM, with supervision from D.V. S.B. handled all mouse immunizations and blood draws. S.B., R.G., and H.S. performed serology experiments.

## SUPPLEMENTARY FIGURES

**Figure S1.**
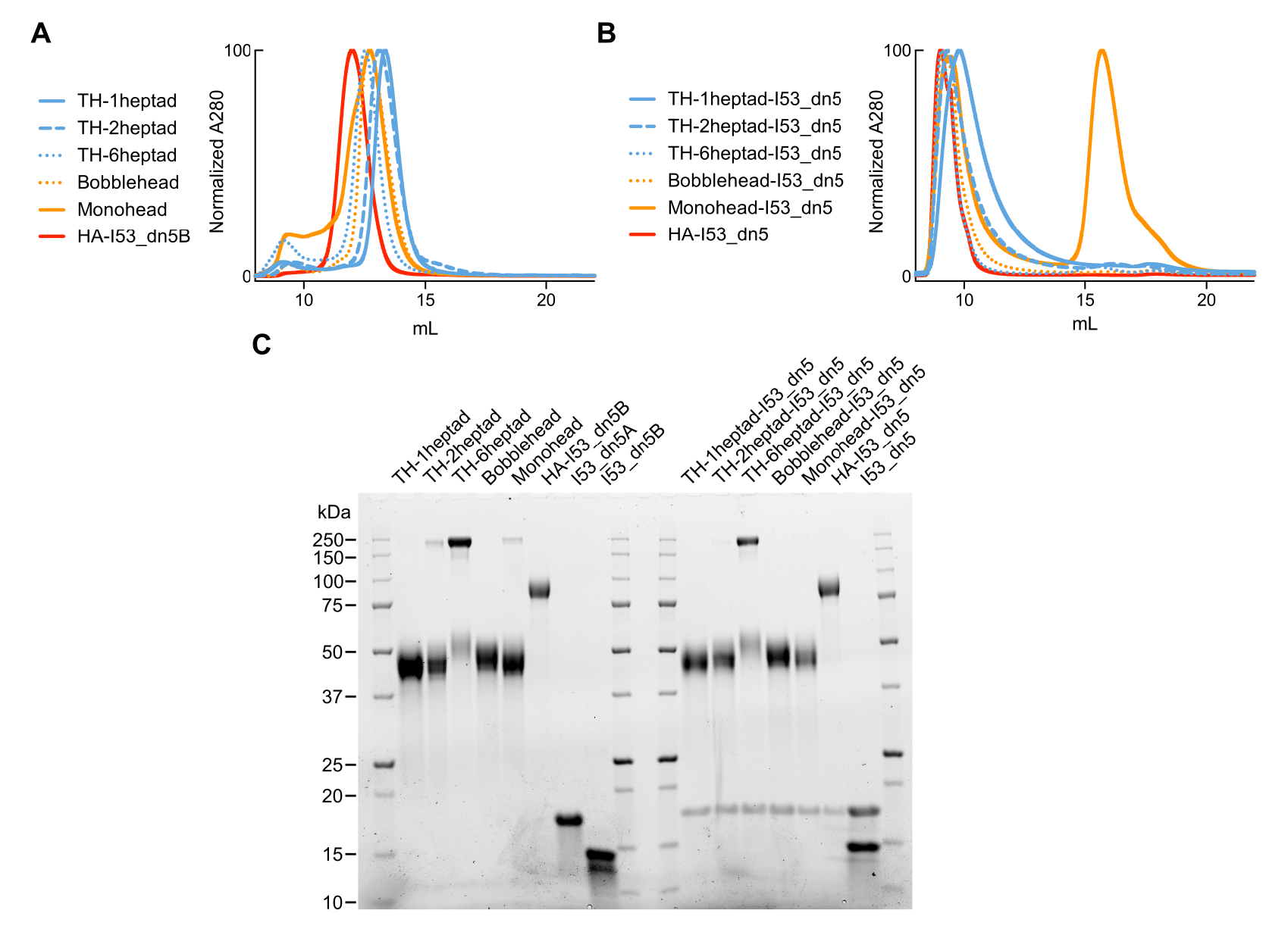
Purification of trihead nanoparticle extension series. (A) SEC chromatogram of trihead extension series components on a Superdex 200 Increase 10/300 GL column. (B) SEC chromatogram of trihead extension series nanoparticles on a Superose 6 Increase 10/300 GL column. (C) SDS-PAGE of trihead extension series components and nanoparticles.

**Figure S2.**
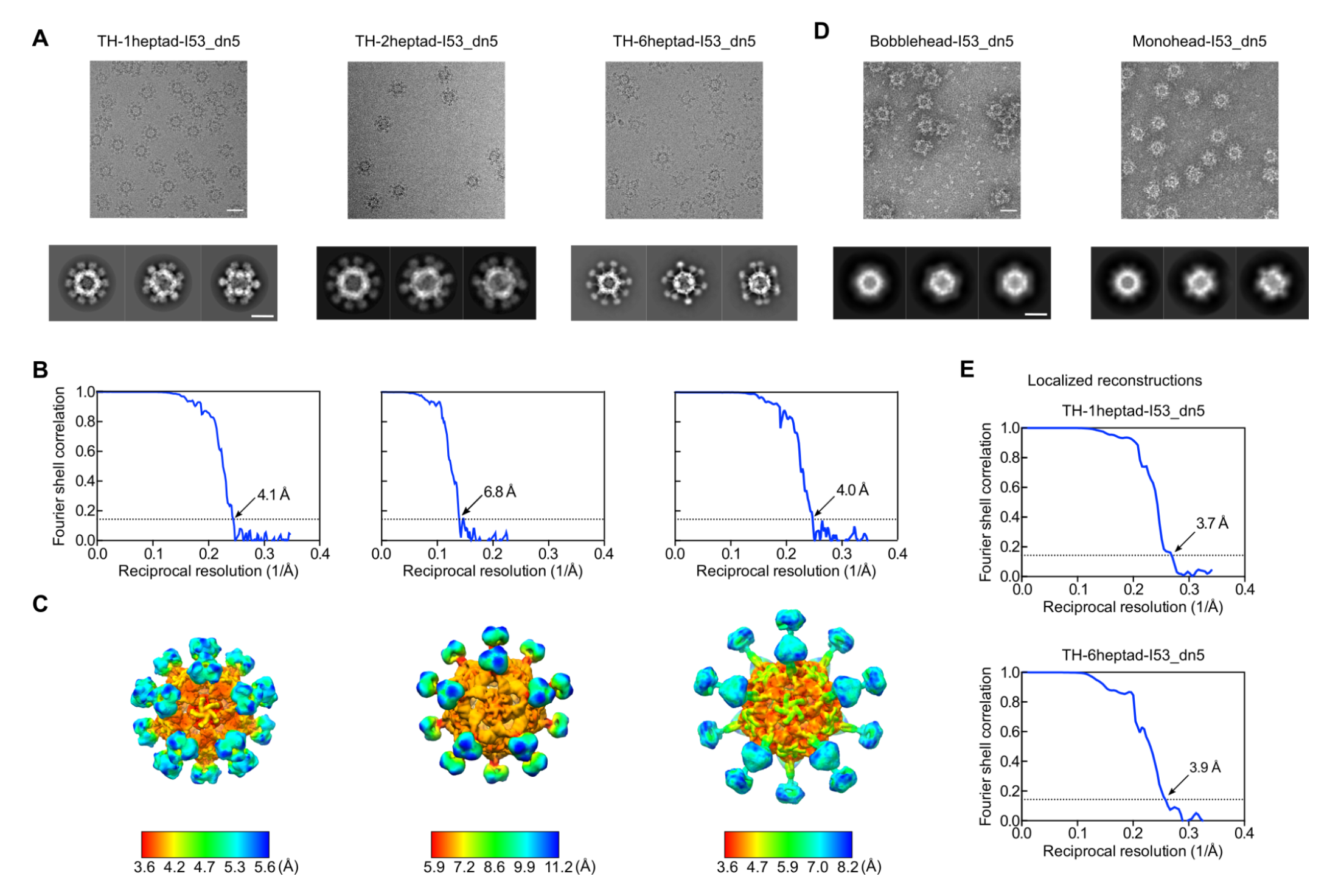
Negative Stain and CryoEM Data Processing, Related to. Figure 3 (A) Representative CryoEM micrographs, scale bar = 50 nm, and 2D class averages, scale bar = 25 nm, of trihead nanoparticles. (B) Gold-standard Fourier shell correlation (FSC) curves for the trihead nanoparticle reconstructions in Figure 3A. The 0.143 cutoff is indicated by the dashed lines. (C) Local resolution maps of the trihead nanoparticle reconstructions. (D) Representative nsEM micrographs, scale bar = 50 nm, and 2D class averages, scale bar = 25 nm, of bobblehead and monohead nanoparticles. (E) Gold-standard FSC curves for TH-1heptad-I53_dn5 and TH-6heptad-I53_dn5 localized reconstructions in Figure 3B.

**Figure S3.**
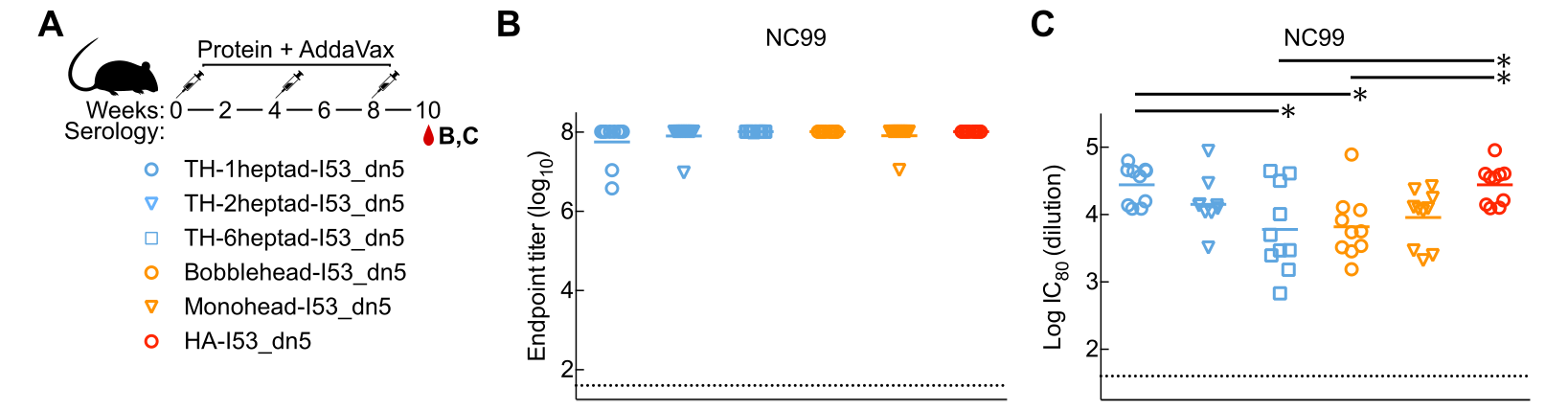
**Immune Responses in Mice Immunized with Trihead Nanoparticle Extension Series** (A) Mouse immunization schedule and groups for adjuvanted trihead nanoparticle extension series. (B) Vaccine-matched NC99 ELISA binding titers in adjuvanted immune sera. (C) Vaccine-matched NC99 ELISA microneutralization titers in adjuvanted immune sera.

**Figure S4.**
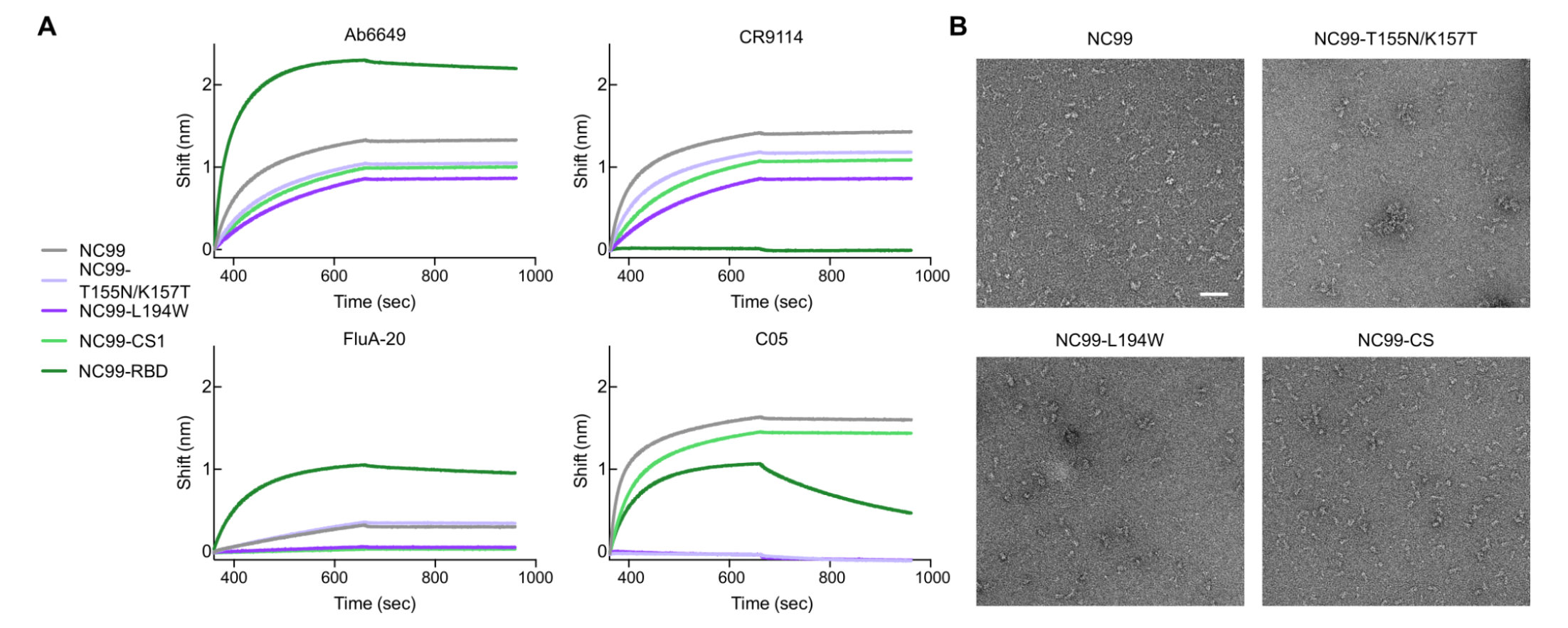
**Biophysical characterization of NC99 probes** (A) BLI of NC99 probes against various HA mAbs. (B) nsEM micrographs of NC99 probes. Scale bar = 50 nm.

**Table S1.**
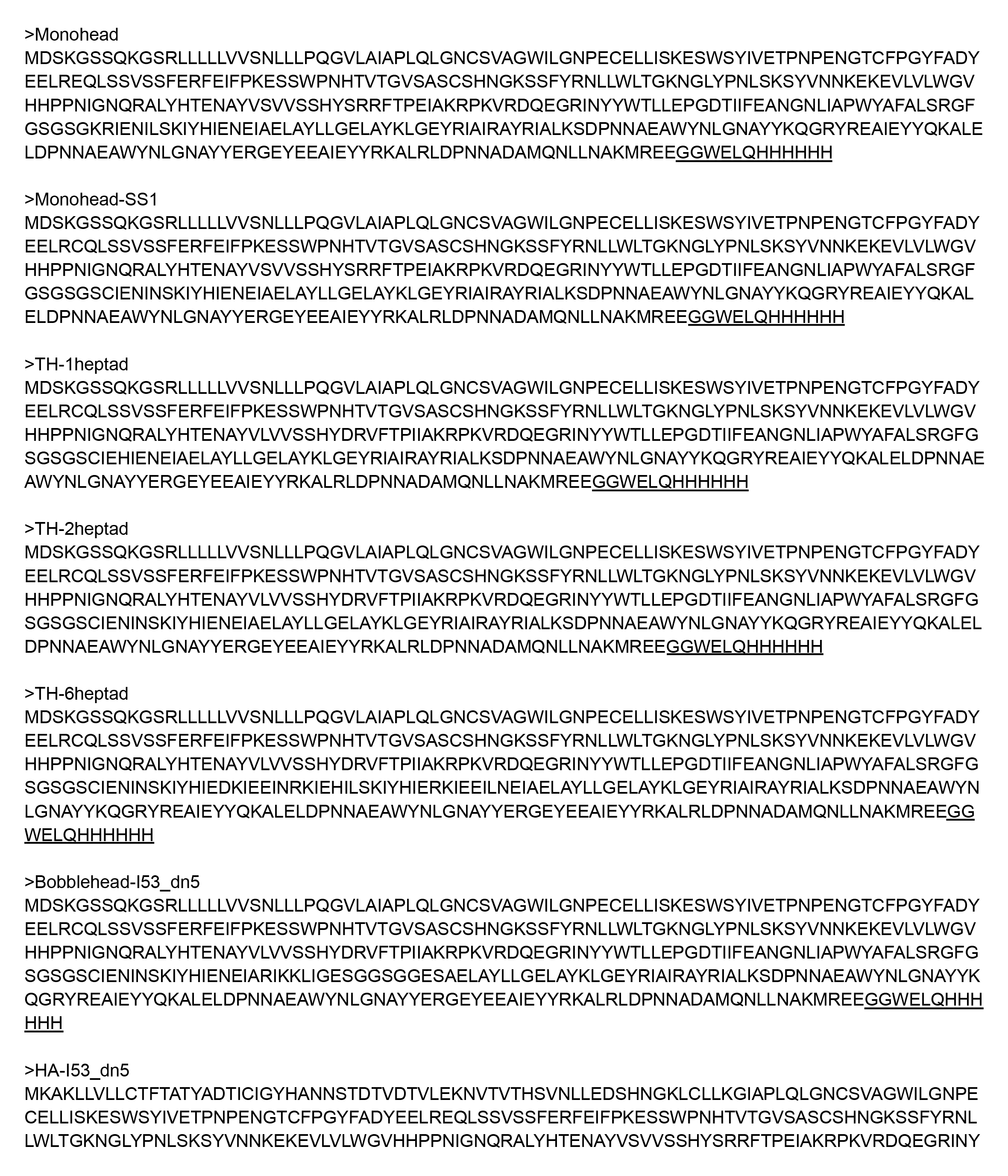
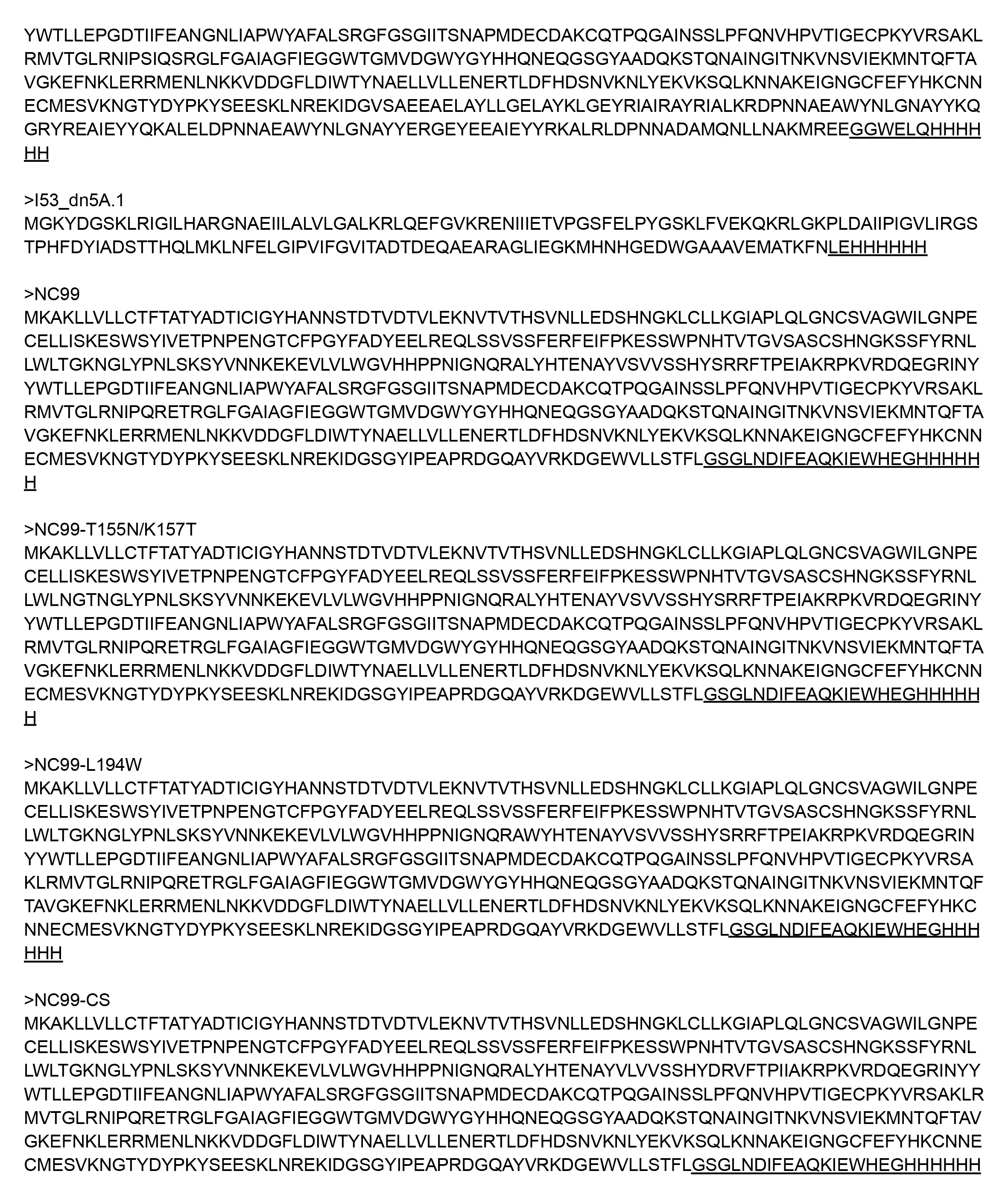
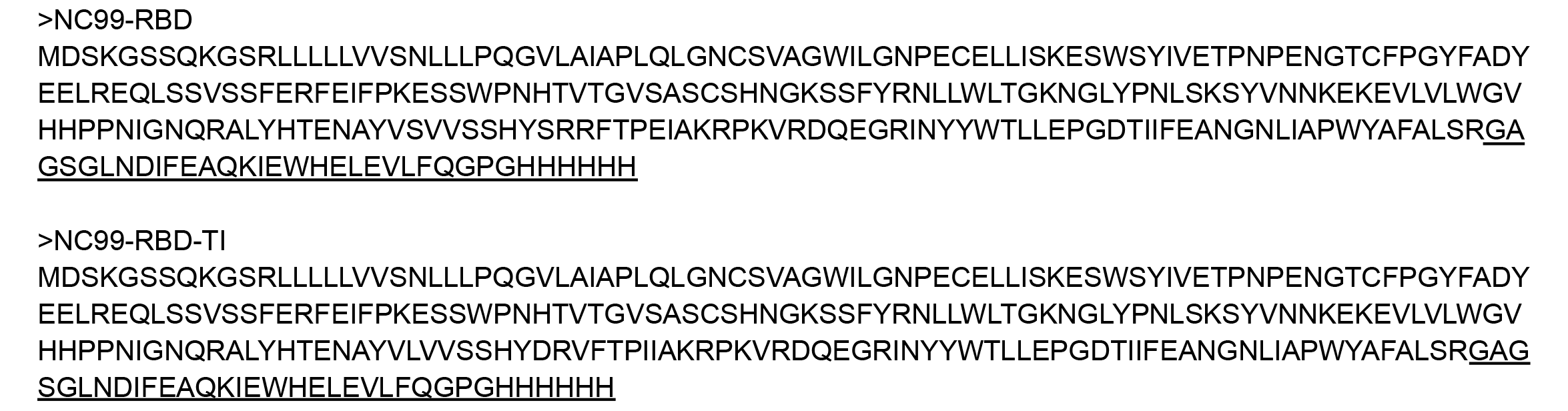
Amino acid sequences of proteins used in this study.

**Table S2.** CryoEM data collection and refinement statistics.

|  | TH-1heptad-I53_dn5<br>EMD XXXX | TH-1heptad-I53_dn5<br>(local refinement)<br>PDB XXXX<br>EMD XXXX | TH-2heptad-I53_dn5<br>EMD XXXX | TH-6heptad-I53_dn5<br>EMD XXXX | TH-6heptad-I53_dn5<br>(local refinement)<br>PDB XXXX<br>EMD XXXX |
| --- | --- | --- | --- | --- | --- |
| **Data collection and processing** |  |  |  |  |  |
| Magnification (×) | 36,000 | 36,000 | 130,000 | 36,000 | 36,000 |
| Voltage (kV) | 200 | 200 | 300 | 200 | 200 |
| Electron exposure (e<sup>-</sup>/Å<sup>2</sup>) | 60 | 60 | 70 | 60 | 60 |
| Defocus range (μm) | -0.5 - -2.5 | -0.5 - -2.5 | -0.5 - -2.5 | -0.5 - -2.5 | -0.5 - -3.5 |
| Pixel size (Å) | 1.16 | 1.16 | 0.525 | 1.16 | 1.16 |
| Symmetry imposed | I | C3 | I | I | C3 |
| Final particle images (no.) | 72,641 | 690,491 | 1,986 | 35,947 | 399,299 |
| Map resolution (Å) | 4.1 | 3.7 | 6.8 | 4.0 | 3.9 |
| FSC threshold | 0.143 | 0.143 | 0.143 | 0.143 | 0.143 |
| Map sharpening B factor (Å<sup>2</sup>) | -215 | -244 | -797 | -186 | -213 |
| **Validation** |  |  |  |  |  |
| MolProbity score |  | 1.86 |  |  | 1.74 |
| Clashscore |  | 6.45 |  |  | 7.52 |
| Poor rotamers (%) |  | 1.71 |  |  | 0.66 |
| Ramachandran plot |  |  |  |  |  |
| Favored (%) |  | 95.22 |  |  | 95.27 |
| Allowed (%) |  | 4.35 |  |  | 4.73 |
| Disallowed (%) |  | 0.43 |  |  | 0 |

